# Disrupted hierarchical organization in disorders of consciousness revealed by fluctuation-dissipation deviations

**DOI:** 10.1101/2025.10.02.679992

**Authors:** Marian Martínez-Marín, Jakub Vohryzek, Anira Escrichs, Dragana Manasova, Jacobo D. Sitt, Morten L. Kringelbach, Yonatan Sanz Perl, Gustavo Deco

## Abstract

Evaluating consciousness levels after coma remains clinically challenging, and probing the brain’s functional hierarchy offers model-based biomarkers of brain states. We characterize the hierarchy loss in disorders of consciousness (DoC) via departures from non-equilibrium dynamics. Irreversible, directed interactions are indexed by deviation from the fluctuation– dissipation theorem (FDT), computed from individualized whole-brain models fit to fMRI from controls and patients in minimally conscious state (MCS) or unresponsive wakefulness syndrome (UWS). Global and resting-state network dynamics in DoC were closer to equilibrium than in controls, decreasing stepwise with decreasing levels of consciousness. Mapping site-specific hierarchical drive over the system revealed disruptions within default-mode network components (e.g., medial and dorsolateral superior frontal gyrus) and subcortical hubs (e.g., thalamus, pallidum and putamen) differentiating between all groups. Recovery of near-control hierarchy in the visual network differentiated MCS from UWS, whereas multiple limbic areas showed similar abnormalities across both DoC groups. Together, these results identify non-equilibrium dynamics as a signature of conscious capacity and stablish FDT deviation as a principled, model-based hierarchy measure that can be operationalised for clinical stratification and monitoring, opening avenues for targeted *in silico* intervention planing.

## Introduction

Evaluating consciousness after brain injury poses significant challenges when relying solely on traditional indicators of arousal and awareness. Signs of arousal (e.g., eye opening) and assessment of awareness based on responsiveness to stimuli depend on voluntary drive and preserved motor output to be evident during examination [1]. The differential diagnosis of minimally conscious (MCS) and unresponsive wakefulness states (UWS) primarily relies on detecting fluctuating yet reproducible signs of awareness that are specific to MCS and appear in addition to the arousal signs observed in UWS. The limitations of conventional diagnostic approaches can lead to misdiagnoses and prognostic inaccuracies in disorders of consciousness (DoC), prompting the development of enhanced diagnostic protocols [1, 2] and the incorporation of neuroimaging-based criteria [3–11].

Despite significant strides in clinical assessment, these approaches may fall short of explaining the pathophysiological mechanisms governing unconscious-state dynamics. In healthy brains, resting-state dynamics are underpinned by a hierarchical, modular organization: network connectivity [12] and hierarchical interactions [13–16] support rich sets of dynamically reconfiguring patterns of neural activity [17]. By contrast, reduced-consciousness states exhibit narrower repertoires of configurations [18–20], and functional connectivity remains closer to anatomical connectivity [21–23]. Computational neuroscience offers diverse model-based frameworks for unpacking the mechanisms behind these observations. Given each individuals’ anatomical connectivity, whole-brain models reproduce functional interactions between nodes lacking direct anatomical links by increasing anatomical fiber conductivity – a global scaling parameter that fits to lower values in DoC relative to controls [24]. Moreover, uncoupling unconscious-like dynamics from explicit anatomical constraints has also been achieved by stimulating specific sites both *in silico* [25, 26] and in non-human primates [27, 28].

Notwithstanding, functional connectivity –statistical dependence between neural signals– does not, on its own, identify causal interactions, which are central to theories of consciousness [29–31]. Model-dependent causal influences between neural masses in the brain are referred to as generative effective connectivity [32], and estimation requires observations that contain dynamical information about directed functional coupling [33]. Cross-covariances can introduce the notion of temporal precedence [34], but predictive influence is typically assessed with Granger-causality-related measures [35]. In empirical perturbation studies, the term effective connectivity often denotes the directed interactions inferred from the propagation of evoked activity after local stimulation [36]. Combining transcranial magnetic stimulation (TMS) with electroencephalography (EEG) has revealed differential breakdown of effective connectivity during unconscious states such as sleep [37, 38], anaesthesia and DoC [4, 39].

Global, dynamics-based indices such as the Perturbational Complexity Index (PCI) capture the spatiotemporal complexity of elicited responses and differentiate levels of consciousness across sleep, anaesthesia and DoC [4]. In computational studies, functional integration has also been used to quantify the brain’s capacity to sustain complex causal interactions under perturbation [40]. Thermodynamics and statistical physics provide principled tools for formalizing the dynamical complexity of neural activity [41, 42]. For spontaneous dynamics, *detailed balance* holds if forward and reverse transition probabilities between configurations are equal; the dynamics are reversible and effectively at equilibrium (Figure 1.a) [43]. DoC dynamics have been classified as closer to equilibrium [44], exhibiting weaker net fluxes within their narrower repertoires of configurations [21, 45–47]. Time-irreversibility can also quantify the direction of information flow between pairs of regions (hierarchical interactions) [35, 48–51], which are closer to reversible (equilibrium) in DoC [52]. Non-equilibrium, irreversible dynamics are a hallmark of asymmetric hierarchical effective connectivity [51, 53, 54].

**Figure 1.**
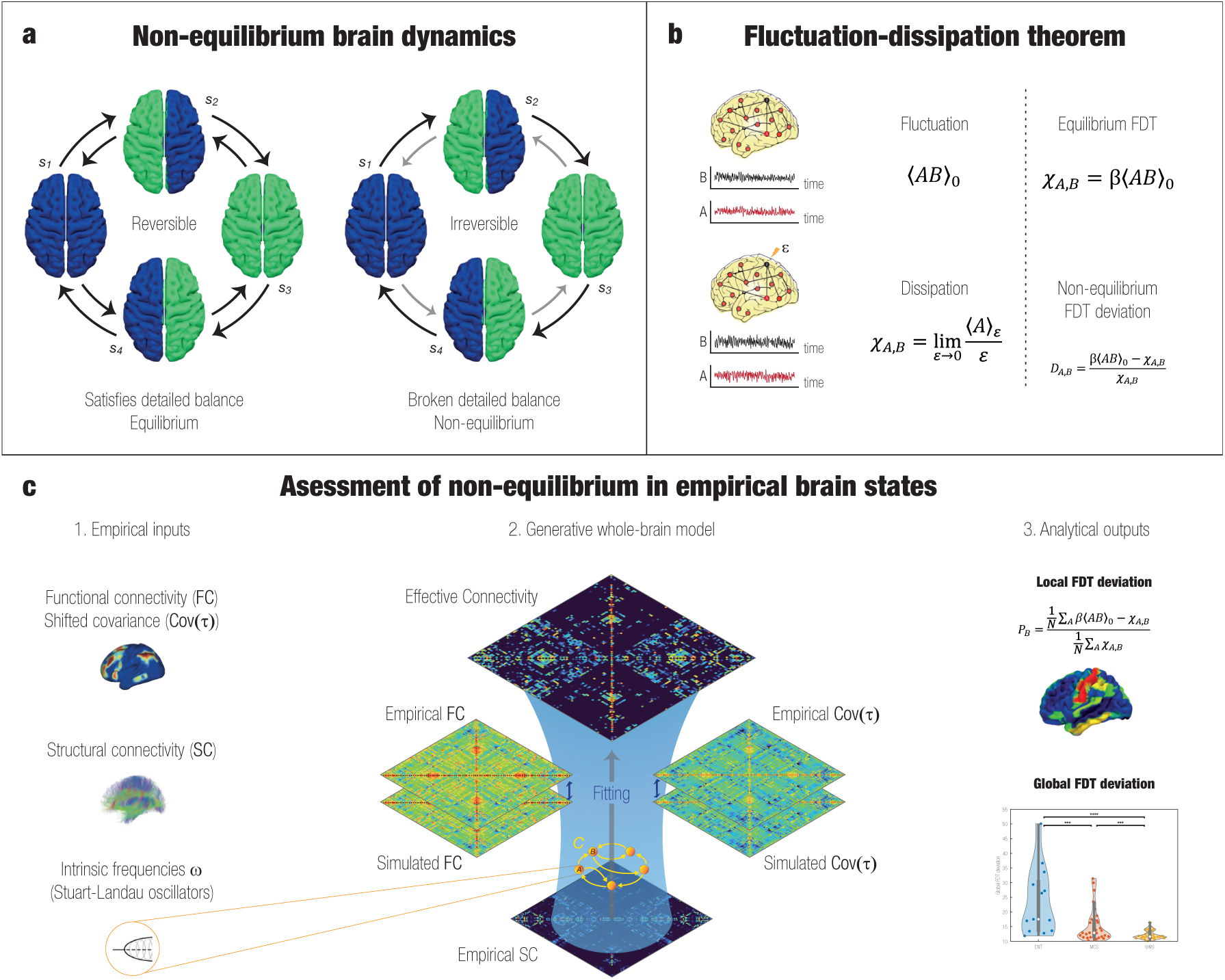
Fluctuation-Dissipation theorem (FDT) deviation as a measure of non-equilibrium in empirical brain states. **a)** Whole-brain activity patterns,*s_i_*, for a limit coarse-grain parcellation (left and right hemispheres) with binarized activity (blue and green) for a given brain state. Transition probabilities (or empirical transition rates) are illustrated by arrows of magnitude-matching intensity. Left: if all the observed transitions are equiprobable, the system satisfies detailed balance and has reversible dynamics. Right: if any pair of transitions between states is not balanced, an arrow of time can be established by the net fluxes of probability. The dynamics are then irreversible and out of equilibrium. Illustration inspired by Fig. 1 from [43]. **b)** Generalization of the FDT to coupled observables: the activity of brain regions *A* and *B*. Left) Correlation between the unperturbed fluctuations and dissipation (the response of *A* after a perturbation *ɛ* is applied on *B*). Right) When the system is initially in equilibrium, the FDT holds and the quantities fluctuation and dissipation are balanced. Deviation from the FDT equilibrium prediction is used as a measure non-equilibrium in the interaction between the ordered pair (*A, B*). **c)** The quantities necessary for computing FDT deviation can be extracted from the equations of coupled Stuart-Landau oscillators that model whole-brain activity. Non-equilibrium dynamics are captured by the asymmetrical effective interactions, fitted to individual empirical neuroimaging data through the combination of anatomical and functional brain connectivity. Local FDT deviation of a region (*B*) is computed as the average deviation across the whole system after its perturbation, and provides a sense of its hierarchical interactions with the system. Average among all local values provides the global FDT deviation, a measure of the overall level of non-equilibrium.

Yet longitudinal DoC studies show that changes in effective connectivity can signal capacity for consciousness before overt changes in ongoing EEG activation become apparent [55]. Here, we analyse closer-to-equilibrium dynamics in DoC using the fluctuation-dissipation theorem (FDT) framework [54], a model-based perturbative approach for quantifying irreversibility in directed interactions. Far from being thought of with the brain as the system of interest, Onsager’s regression principle states that –for a system initially in equilibrium– spontaneous fluctuations of an observable predict its linear response to a weak perturbation [41] (Figure 1.b). Deviations from this prediction provide a measure of non-equilibrium [56]. Its extension to coupled observables enables the quantification of non-equilibrium directed interactions by comparing equilibrium prediction with the actual model-based response to site-specific perturbations [54]. After fitting individualised Hopf whole-brain models, we exploit closed-form expressions of linear response to compute the causal response of every region to infinitesimal, site-specific perturbations (Figure 1.c). The mismatch between this response and the equilibrium prediction –FDT deviation– indexes broken detailed balance, time-irreversibility, and hierarchical effective coupling. This yields a principled, model-based measure that can be mapped at global, network, and regional scales. We assess the degree of non-equilibrium –globally and within resting-state networks– to differentiate the brain states underlying the acquired neuroimaging data. A whole-brain functional hierarchy is obtained by aggregating each region’s downstream causal impact. Regional hierarchical levels are compared between conditions, with a special focus on potential recovery-related similarities and differences between MCS and UWS.

## Methods

### Participants

A total of 13 healthy controls (7 females, mean age ± SD, 42.54 ± 13.64 years) and 53 patients with DoC were included in this study, previously described in [52, 57, 58]. Patients were hospitalised in Paris Ĥopital Pitié-Salp^etrìere after brain injury. Clinical assessment and trained clinicians evaluated their Coma Recovery Scale-Revised (CRS-R) scoring to determine the state of consciousness. Patients were diagnosed with UWS if they showed arousal (eye-opening) and no signs of awareness (non-reflex voluntary movement). Conversely, those exhibiting behaviours indicative of awareness (e.g., visual pursuit, response to pain or reproducible command following) were diagnosed with MCS. Four MCS patients from the original dataset, presenting null signal in several regions, were excluded. The provided sex and age information for this group referred to the original 33-patients sample. Everything considered, DoC groups in this study included 29 MCS patients (7-11 females, mean age ± SD, 47.2±20.76) and 24 UWS patients (9 females, mean age ± SD, 39.25±16.30 years). This research was approved by the local ethics committee Comité de Prection des Personnes Ile de France 1 (Paris, France) under the code ‘Recherche en soins courants’ (NEURODOC protocol, n° 2013-A01385-40). The patient’s family grave their informed consent for the participation of their relative, and all investigations were conducted according to the Declaration of Helsinki and French regulations.

### MRI data acquisition

Participants’ fMRI data was acquired with two different acquisition protocols. In the first, 13 healthy controls and 21 patients (11 MCS, 10 UWS) were scanned on a 3T General Electric Signa System. Resting-state fMRI images were acquired axially with a gradient-echo EPI sequence (200 volumes, 48 slices of thickness 3mm, TR/TE: 2400ms/30ms, voxel size 3.4375×3.4375×3.4375 (mm)^3^, flip angle 90°, FOV 220mm^2^). In the second protocol, 34 patients (20 MCS, 14 UWS) were scanned on a 3T Siemens Skyra System. Resting-state fMRI images were acquired axially with a gradient-echo EPI sequence (190 volumes, 62 slices of thickness 2.5mm, TR/TE= 2000ms/30ms, voxel size 2×2×2 (mm)^3^, flip angle 90°, FOV 220 mm^2^, multiband factor 2).

Generative effective connectivities were initialized using the anatomical connectivity matrix described in [59], from Human Connectome Project (HCP) diffusion- and T2-weighted imaging data of 32 unrelated healthy subjects (14 females, mean age ± SD, 31.5 ± 8.6 years; scanned for 89 minutes each). The dataset and acquisition details are available in the Image & Data Archive under HCP project (https://ida.loni.usc.edu/). The preprocessed data is part of the freely available Lead-DBS software package (http://www.lead-dbs.org/). Additionally, data from 100 unrelated HCP subjects (U100) was used to compute the correaltion between AAL subcortical voxels and Yeo’s resting-state networks cortical voxels [60]. Specifically, the volumetric timeseries *rfMRI REST1 LR hp2000 clean.nii.gz*, with acquisition and denoising protocols detailed in [61]. Briefly, a customized Siemens 3T “Connetome Skyra” scanner was used to acquire oblique axial LR acquisition with a gradient-echo EPI sequence, 73 slices of thickness 2mm, TR/TE:720ms/33.1ms, 2.00 isotropic voxels, flip angle 52 deg, FOV 208×180mm^2^). Further information can be found on the HCP website (http://www.humanconnectome.org/).

### fMRI preprocessing

All data was preprocessed using FSL (http://fsl.fmrib.ox.ac.uk/fsl) as described in our previous study [24]. Resting-state fMRI was computed using MELODIC (Multivariate Exploratory Linear Optimised Decomposition into Independent Components) [62]. The steps included exclusion of the first five volumes, MCFLIRT for motion correction [63], Brain Extraction Tool (BET) [64], spatial smoothing with 5 mm FWHM Gaussian Kernel, rigid-body registration, high pass filter cutoff=100.0s, and single-session ICA with automatic dimensionality estimation. Additionally, noise components and patient’s lesion-related artifacts were regressed out independently for each subject using FIX (FMRIB’s ICA-based X-noiseifier) [65]. Finally, FSL tools were used for individual’s image co-registration and extraction of the time-series of *N* = 90 cortical and subcortical brain areas from MNI space using Automated Anatomical Parcellation (AAL) [66].

### Probabilistic tractography analysis

An anatomical connectivity matrix, previously used in [59], was derived from diffusion- and T2-weighted MRI. Diffusion data was processed using a generalized q-sampling imaging algorithm via DSI studio (http://dsi-studio.labsolver.org/). A white-matter mask, generated from the segmentation of the T2-weighted anatomical images, was used for co-registration of the images to the b0 images of the diffusion data using the SPM12 toolbox (https://www.fil.ion.ucl.ac.uk/ spm/software/spm12/). For each participant, the 200,000 most probable fibers within this mask were sampled and transformed to the Montreal Neurological Institute (MNI) space. Individual tractograms were aggregated in MNI standard space, forming a normative tractogram (accessible through the leadDBS toolbox) from which the matrix was computed with AAL parcellation scheme.

### Inclusion of subcortical regions in resting-state networks

Subcortical regions included in AAL parcellation were related to Yeo’s 7 resting-state networks [67] using resting-state volumetric timeseries from 100 unrelated HCP subjects (U100) [61] (isotropic 2mm voxels). Following Tian *et al*., a spatial Gaussian smoothing with FWHM = 6mm was applied to each time frame, followed by Wishart filtering denoising with their function *icaDim* [68, 69]. Following [67], a 0.008-0.08 Hz butterworth filter was applied.

Subcortical voxels were labelled after AAL [70], while cortical ones were assigned both Schaefer 1000 and Yeo 7 labels [67]. Individual functional connectivities between subcortical and cortical voxels were computed. Given the group-averaged functional connectivity, each subcortical voxel obtained an average correlation per resting-state network, and was associated to the network with maximum average correlation (Figure 3.a; Supplementary Figure 6 and Table 1).

**Table 1:** Subcortical RSN assignment using 100 unrelated HCP subjects. For each AAL subcortical ROI (L/R), the table reports the percentage of voxels assigned to each Yeo 7 network and 95% group–bootstrap percentile CIs. Resampling was at the subject level (*B* = 10,000, fixed seed); for each resample we recomputed the group-averaged subcortical–cortical FC, reassigned each subcortical voxel to the RSN with maximal mean FC (argmax), and tallied voxel percentages within the ROI. CIs are percentile. Preprocessing followed [67–69].

| Label | Region | Visual |  | Somato. |  | D. Att. |  | V. Att. |  |
| --- | --- | --- | --- | --- | --- | --- | --- | --- | --- |
|  |  | % | CI | % | CI | % | CI | % | CI |
| 38 | L Hippocampus | 33.2 | [23.5, 42.4] | 54.1 | [44.9, 61.4] | 9.4 | [5.6, 14.3] | 0.0 | [0, 0.1] |
| 37 | R Hippocampus | 26.4 | [17.7, 33.6] | 67.3 | [59.2, 74.4] | 2.5 | [1, 6] | 0.0 | [0, 0.2] |
| 42 | L Amygdala | 1.6 | [0.6, 8.5] | 98.0 | [89.5, 99.2] | 0.0 | [0, 0.8] | 0.4 | [0, 4.4] |
| 41 | R Amygdala | 0.0 | [0, 0.9] | 99.1 | [94.1, 100] | 0.0 | [0, 0] | 0.5 | [0, 5] |
| 72 | L Caudate | 12.7 | [7.6, 21.9] | 0.3 | [0.1, 5.1] | 1.0 | [0.2, 7] | 19.3 | [4.9, 22.5] |
| 71 | R Caudate | 8.0 | [3.1, 16.3] | 2.8 | [0.7, 12.4] | 1.0 | [0, 7.2] | 2.6 | [0, 5.5] |
| 74 | L Putamen | 2.6 | [0.6, 7.9] | 28.2 | [23.9, 42.8] | 1.2 | [0.4, 5.9] | 67.1 | [48, 68.9] |
| 73 | R Putamen | 0.5 | [0, 6.1] | 9.4 | [5.6, 24.3] | 0.0 | [0, 3] | 86.6 | [66, 86.9] |
| 76 | L Pallidum | 0.4 | [0, 7.1] | 23.2 | [15.4, 46.1] | 0.0 | [0, 8.2] | 74.6 | [45.4, 77.1] |
| 75 | R Pallidum | 10.6 | [2.7, 24.6] | 3.8 | [0.3, 28.2] | 2.0 | [0, 16] | 71.7 | [38.6, 74.1] |
| 78 | L Thalamus | 16.6 | [7.6, 27.5] | 45.3 | [35.6, 55.1] | 11.4 | [7.2, 17.9] | 3.5 | [0.2, 7.8] |
| 77 | R Thalamus | 4.8 | [1.6, 18] | 44.5 | [34.5, 52.6] | 11.1 | [5.3, 19.1] | 7.0 | [0.5, 10] |

| Label | Region | Limbic |  | Control |  | DMN |  |
| --- | --- | --- | --- | --- | --- | --- | --- |
|  |  | % | CI | % | CI | % | CI |
| 38 | L Hippocampus | 3.0 | [1.1, 5.1] | 0.0 | [0, 1.2] | 0.3 | [0.1, 4.1] |
| 37 | R Hippocampus | 1.2 | [0.2, 2.9] | 0.1 | [0, 1.2] | 2.6 | [1.4, 5.3] |
| 42 | L Amygdala | 0.0 | [0, 0] | 0.0 | [0, 0] | 0.0 | [0, 0] |
| 41 | R Amygdala | 0.0 | [0, 0] | 0.0 | [0, 0] | 0.5 | [0, 0.9] |
| 72 | L Caudate | 33.6 | [23.4, 49.3] | 0.0 | [0, 0.1] | 33.1 | [24.6, 42.3] |
| 71 | R Caudate | 34.5 | [23.6, 44] | 0.0 | [0, 0.1] | 51.0 | [34.9, 61.3] |
| 74 | L Putamen | 0.0 | [0, 2.8] | 0.0 | [0, 0] | 0.8 | [0, 5] |
| 73 | R Putamen | 1.1 | [0.5, 12.4] | 0.0 | [0, 0] | 2.4 | [1.1, 5.2] |
| 76 | L Pallidum | 0.0 | [0, 8.9] | 0.0 | [0, 0] | 1.8 | [0, 4.6] |
| 75 | R Pallidum | 5.8 | [0.7, 20.1] | 0.0 | [0, 0] | 6.1 | [1.7, 14.3] |
| 78 | L Thalamus | 13.8 | [7.4, 21.8] | 0.0 | [0, 0] | 9.5 | [5, 13.9] |
| 77 | R Thalamus | 25.1 | [15.5, 34.3] | 0.0 | [0, 0] | 7.5 | [3.7, 13.7] |

### Fluctuation-Dissipation theorem

As put forward in [54], considering the activity of each pair of regions as coupled observables, *A*(*t*) and *B*(*t*), the asymmetry in the interaction B-to-A can be quantified by applying a sub-threshold perturbation *ɛ* to *B*(*t*) at *t* = 0 and observing how deviated from the equilibrium prediction is the response in *A*(*t*). The difference between the expected value of *A* in absence of the perturbation, ⟨*A*(*t*)⟩_0_, and after the perturbation, ⟨*A*(*t*)⟩*_ɛ_*, can be expressed in terms of the simultaneous and lagged co-behaviour as

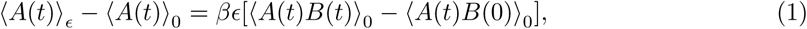

with *β* the inverse temperature. Susceptibility to the perturbation is defined as

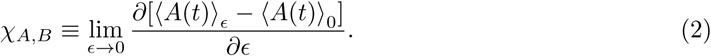

According to fluctuation-dissipation theorem, the static susceptibility in equilibrium is given by the unperturbed correlation

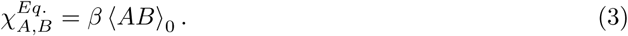

For non-equilibrium dynamics, comparing Equation 3 to the actual linear response to the perturbation (*χ_A,B_*, Equation 12) provides a measure of irreversibility in the B-to-A interaction. The hierarchical drive from region B to A is quantified by the FDT deviation, defined as

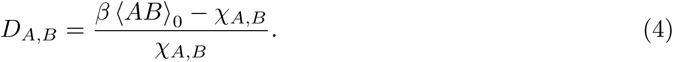

### Whole-brain model

For each pair of brain regions (*i, j*), the difference between the equilibrium prediction and the actual response of *i* when a weak external perturbation is applied onto *j* (Equation 4) is calculated *in silico* using a whole-brain generative model. For each brain region, the Hopf whole-brain model [71] describes the activity of an isolated node *j* with a Stuart-Landau oscillator,

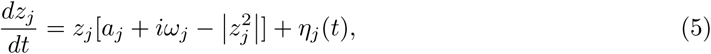

where *z_j_* = *x_j_* + *iy_j_* is a complex number whose real part, *Re*(*z_j_*) = *x_j_* describes the BOLD signal, *a_j_* is the bifurcation parameter, *ω_j_* is the intrinsic frequency of each region, and *η_j_*(*t*) additive Gaussian noise with standard deviation *σ*. The associated *N* equations were coupled by means of a generative effective connectivity matrix **C** under *common difference coupling*. The activity of the *j*-th node in a coupled network with *N* oscillators is then given by

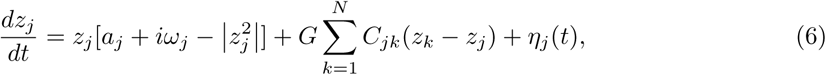

with a global parameter *G* scaling the conductivity for all connections. Expressing the activity of all nodes as a vector **z** = [*z*_1_*, …, z_N_*]*^T^*,

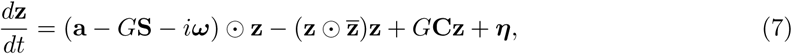

with vectors **a** = [*a*_1_*, …, a_N_*]*^T^*, ***ω*** = [*ω*_1_*, …, ω_N_*]*^T^*, ***η*** = [*η*_1_*, …, η_N_*]*^T^*, and **S** for the bifurcation parameters (*a_j_*=-0.02), the intrinsic frequencies (estimated as the averaged peak frequency of the narrowband BOLD signals), additive uncorrelated Gaussian noise (standard deviation *σ* = 0.01), and the connectivity strength of each node (*S_i_* = ∑*_j_ C_ij_*). The superscript [*…*]*^T^* represents the transpose vector, **z̄** is the complex conjugate of **z**, and ⊙ is the Hadamard element-wise product.

Following [54, 72], the evolution of the fluctuations around the fixed point **z=0** of Equation 7 can be approximated linearly by

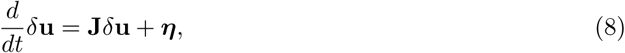

where the 2*N* -dimensional vector *δ***u** = [*δ***x***, δ***y**]*^T^* = [*δx*_1_*, …, δx_N_, δy*_1_*, …, δy_N_*]*^T^* contains the fluctuations of the real and imaginary state variables. The Jacobian matrix **J** can be written as a block matrix

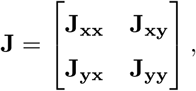

with *N* × *N* matrices **J_xx_** = **J_yy_** = *diag*(**a** − *G***S**) + *G***C** and **J_xy_** = −**J_yx_** = *diag*(***ω***), being *diag*(**v**) the matrix with diagonal values given by the vector **v**. Equation 8 allows an analytical solution for the simulated covariance matrix **K** = 〈*δ***u***δ***u***^T^*〉, which is necessary for quantifying deviation from equilibrium (Equation 4), if rewritten as a Sylvester equation

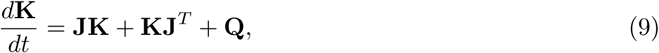

with a noise covariance matrix that is diagonal for uncorrelated noise, **Q** = *diag*(*σ*^2^). The stationary covariance 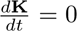 can be solved analytically (`MATLAB: sylvester`), and the simulated functional connectivity **FC***^model^*can be derived from its first *N* rows and columns (`MATLAB: corrcov`).

This model was fitted individually to the BOLD fMRI empirical data for each subject by optimizing the effective connectivity matrix, **C**, and the global coupling parameter *G*.

For *C* to capture empirical non-equilibrium dynamics, asymmetrical constrains given by the lagged correlation **KS***^empirical^* (*τ* =, normalising 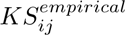 by 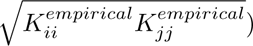) were considered along with the symmetric ones provided by the anatomical (**SC**) and functional (**FC***^empirical^*) connectivities [34, 54]. The normalised simulated shifted covariance is calculated as **KS***^model^* = *e^τ^***^J^K**, prior **K** normalisation dividing *K_ij_* by 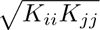. A pseudo-gradient descent procedure was used to update **C**, following

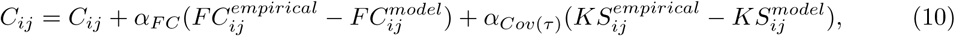

with *α_F C_* = *α_Cov_*_(*τ*)_ = 0.0005. First, group’s **C***_group_* were initialized from **SC** and iteratively compared to group-averaged 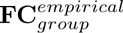 and 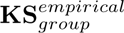. Then, individual **C** were initialized with their **C***_group_* and optimized for the subject’s **FC***^empirical^*and **KS***^empirical^*. Finally, each subject’s global coupling parameter *G* was selected as the one between 0.5 to 1.5 (step size 0.01) minimizing the mean squared errors between the empirical an simulated **FC** and **KS** matrices.

### Deviation from equilibrium

A perturbation in node *j* can be introduced to the evolution of fluctuations (Equation 8) with a 2*N* -dimensional vector, **h**_(*j*)_, where *h*_(*j*)*i*≠*j*_ = 0 and *h*_(*j*)*i*=*j*_ = 1 [54]. Taking the statistical average,

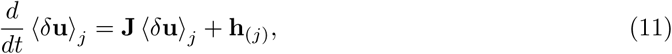

the stationary linear response follows as

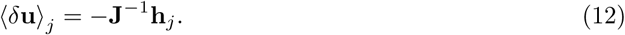

The real part ⟨*δ***x**⟩*_j_* allows computing FDT deviation for region *i* after perturbing *j* as

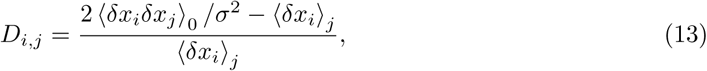

where 2*/σ*^2^ acts as the inverse temperature *β* and 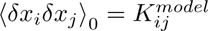. Per perturbation site *j*, its local FDT deviation is the aggregation of non-equilibrium responses over measuring sites *i*:

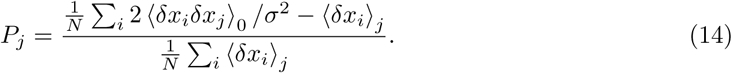

For each brain region, *P_j_* quantifies the average non-equilibrium in its (directed) interactions with other regions (Figure 4.a), with high values indicating more hierarchical interactions [41]. The whole-brain average level of non-equilibrium, referred to as global deviation (Figure 2.a), was computed as the average local FDT deviation over all perturbation sites 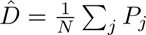. The coefficient of variation in FDT deviation was used for measuring hierarchical diversity (Figure 2.b).

**Figure 2.**
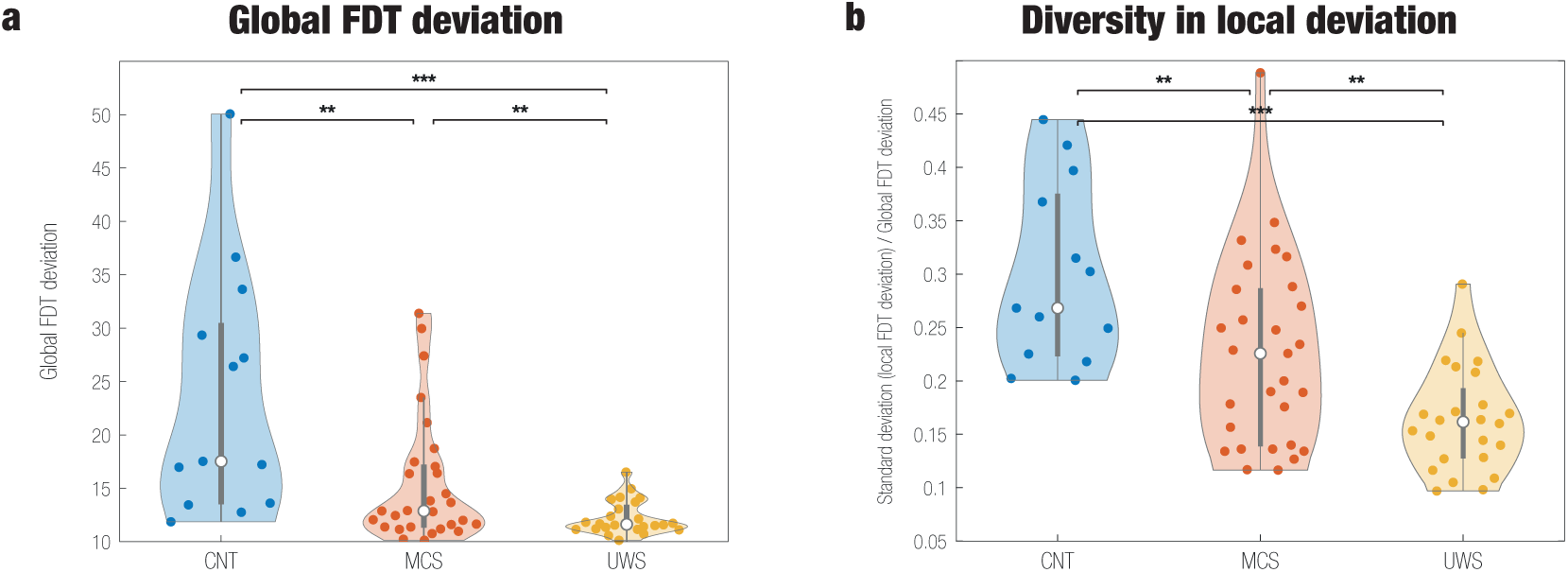
Global FDT deviation and hierarchical diversity in the brain. **a) Global FDT deviation.** Each subject’s average local FDT deviation across the whole brain is represented (dots). Pairwise group comparisons show smaller global deviation for less conscious states, indicative of closer to equilibrium dynamics. **b) Hierarchical diversity across the whole brain.** The coefficient of variation of FDT deviation per subject (standard deviation among local FDT deviations normalized by the average deviation), computed as an indicator of hierarchical diversity across the whole brain. In all comparisons, decreasing levels of consciousness exhibit less hierarchical diversity (* *p* < 0.05, ** *p* < 0.01, *** *p <* 0.001).

At the level of the seven resting-state networks defined by Yeo *et al*. (2011) [67], the average deviation per network was computed as a volume-weighted average due to the distributed network-membership of AAL regions [70]. Specifically, a map with the number of 2mm^3^ MNI voxels per AAL region that belong to each RSN was used. For *v_RSN,j_* tne number of voxels of each region *j* in a RSN, the network average deviation follows as

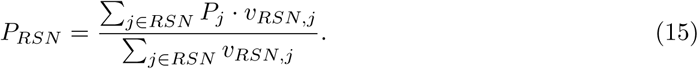

The weighted local deviations, *P_j_* · *v_RSN,j_/* ∑*_j_*_∈_*_RSN_ v_RSN,j_*, are referred to as *contributions*. Subcortical regions (hippocampus, amygdala, caudate nucleus, putamen, pallidum, and thalamus) were previously voxel-wise associated to one of the networks based on the maximum average functional correlation to the respective cortical voxels (Figure 3.a). Hierarchical diversity within each network was computed as the coefficient of variation of the contributions (the standard deviation of *P_j_* · *v_RSN,j_/* ∑*_j_*_∈_*_RSN_ v_RSN,j_* normalized by the average network deviation *P_RSN_*, Figure 3.b).

**Figure 3.**
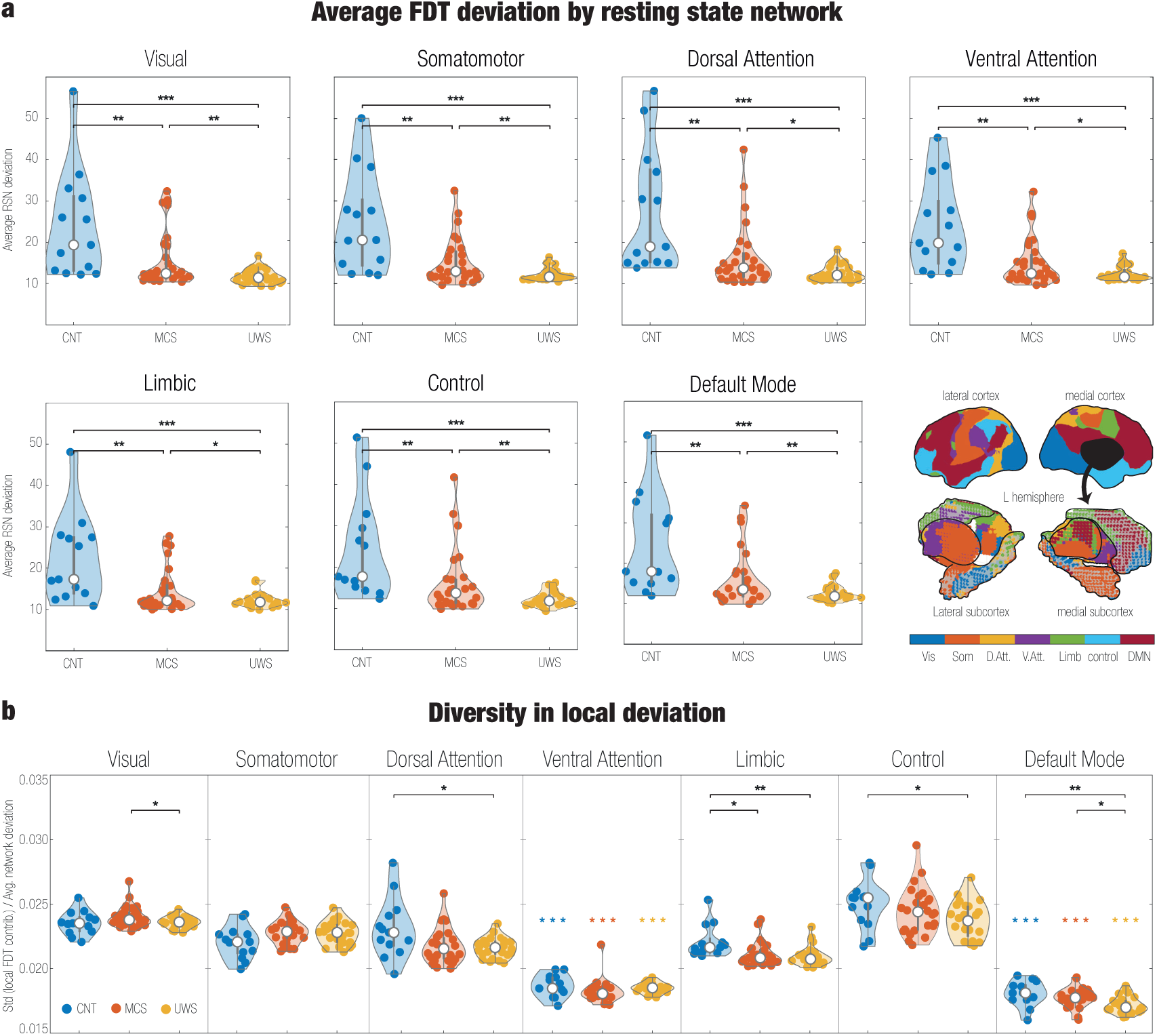
FDT deviation by resting-state network and hierarchical diversity among their components. **a) Average FDT deviation within resting-state and subcortical networks.** Average FDT deviation for each of the seven resting-state networks defined by Yeo *et al*. [67] (visual, somatomotor, dorsal attention, ventral attention, limbic, control and default mode networks). Subcortical regions were included in the networks averages (see Methods §). Low network average deviation indicates that the interactions of its components toward the whole-brain system are closer to equilibrium. Across all networks and group pairwise comparisons, less conscious states present smaller network average deviation (FDR corrected per pair of groups, *N* = 7). **b) Hierarchical diversity within resting-state networks**. For each subject, the coefficient of variation of FDT deviation within the networks was computed as the standard deviation of the (weighted) local FDT deviations normalized by the network’s average deviation (see Methods). The coefficient of variation in local deviation within the networks is an order of magnitude smaller than that across the whole brain (Figure 2.b). Compared to controls, both MCS and UWS presented significant differences in within-network hierarchical diversity for all networks except for the visual. Among these differences, a tendency for decreased diversity was observed for all except for the somatomotor network. Between the MCS and UWS groups, none of the distributions of hierarchical diversity per network differed significantly (FDR corrected per pair of groups, *N* = 7). Additionally, hierarchical diversity across networks per group was significantly smaller in the default mode regarding other networks, followed by the ventral attention (FDR corrected per group, *N* = 21; * *p <* 0.05, ** *p <* 0.01, *** *p <* 0.001).

### Statistical analyses

Comparison of the FDT deviation values (global, RSN and local) between each pair of groups was performed using a non-parametric permutation-based unpaired t-test. After 5000 permutations of group labels, the null distribution is estimated independently for each group. For each pair of groups, comparisons of network average deviation (Figure 3.a), hierarchical diversity within networks (Figure 3.b), and local deviation (Figure 4.a) were corrected for multiple comparisons with Benjamini–Hochberg False Discovery Rate (FDR, *N* = 7, 7, 90 respectively), considering p-value significance below 0.05 [73].

**Figure 4.**
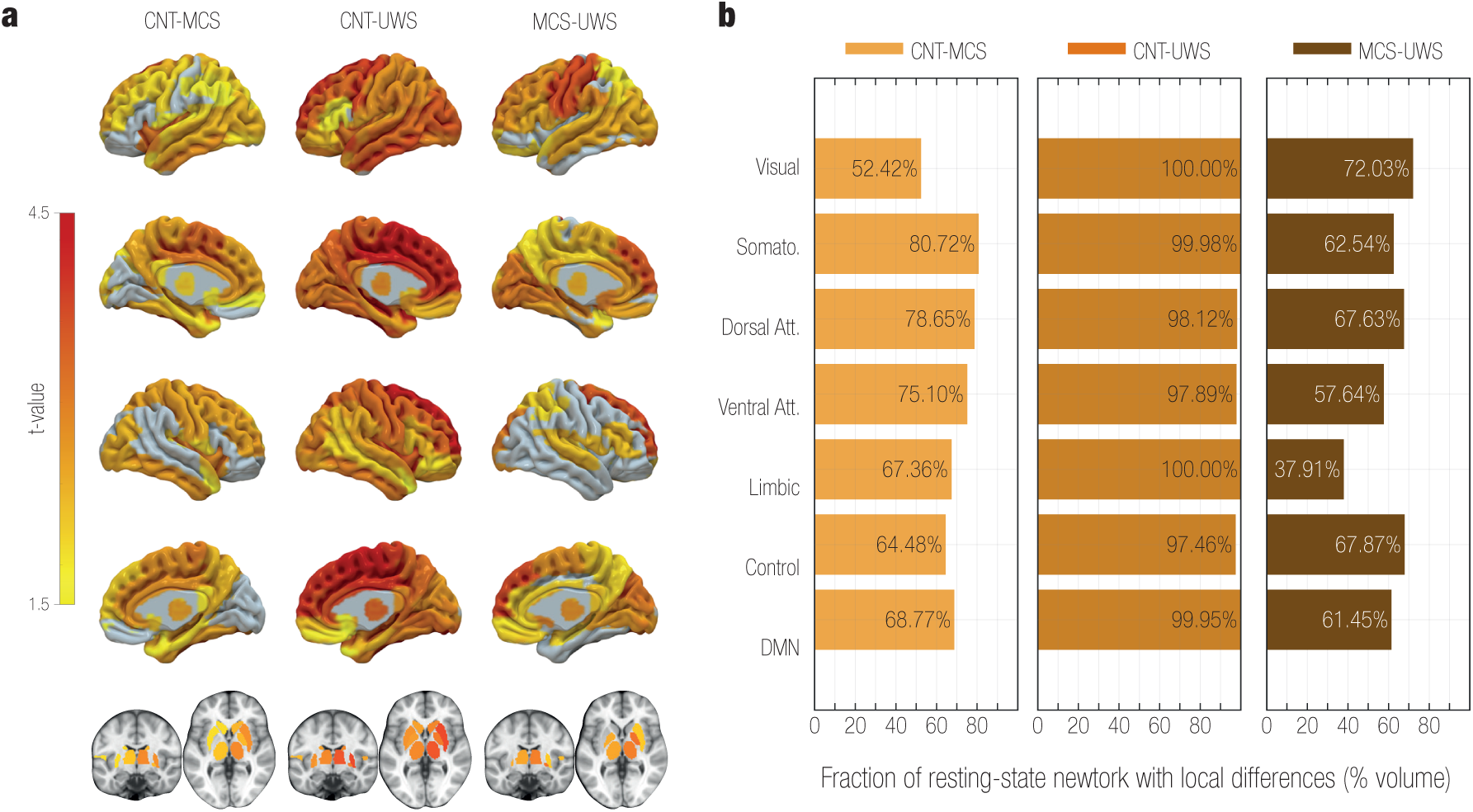
Local FDT deviation. **a) Differences in local FDT deviation between groups.** For each pairwise group comparison (columns), all significant t-values (colour scale; FDR corrected, *N* = 90) were positive, indicating greater local deviations for the first group. Gray areas did not show significant differences. **b) Fraction of resting-state network involved in the local differences.** For each pairwise group comparison (coloured labels), the plot shows the RSN volume fraction occupied by the subsets of local differences in a). Local differences underpin the global and RSN averaged results in Figures 2.a and 3.a. Although all global and RSN averages differed across groups, indicating that local differences sufficed to impact the averages, distinct amounts of each network were involved in such differences.

## Results

### Global level of non-equilibrium

The FDT framework was applied to resting-state control (CNT), MCS and UWS groups. Global FDT deviation was computed as the average local deviation across the whole brain (Equation 14). Global deviation decreased stepwise with level of consciousness (Figure 2.a; *p_CNT_* _−*MCS*_ *<* 0.01*, p_CNT_* _−*UW*_ *_S_ <* 0.001*, p_MCS_*_−*UW*_ *_S_ <* 0.01; FDR-corrected), indicating increasingly closer to equilibrium dynamics in less conscious states. We next quantified hierarchical diversity as the coefficient of variation of local deviations across regions. Hierarchical diversity likewise declined with decreasing consciousness (Figure 2.b; *p_CNT_* _−*MCS*_ *<* 0.01*, p_CNT_* _−*UW*_ *_S_ <* 0.001*, p_MCS_*_−*UW*_ *_S_ <* 0.01), consistent with a flattening of functional hierarchy in DoC.

### Resting-state network average level of non-equilibrium

For the seven resting-state networks (RSNs) defined by Yeo *et al*. –visual, somatomotor, dorsal attention, ventral attention (V.Att.), limbic, control and default mode (DMN)–, we computed volume-weighted, network-average FDT deviation (Equation 15). Subcortical voxels were assigned to the cortical RSN with maximal average correlation derived from 100 unrelated HCP subjects (see Methods). All networks exhibited progressively closer-to-equilibrium dynamics with decreasing consciousness (Figure 3.a; FDR-corrected).

Within-network hierarchical diversity (coefficient of variation of local deviations) was small relative to whole-brain diversity (compare the y-axis from Figure 3.b and Figure 2.b). Within each group, the default mode and ventral attention networks showed the lowest diversity (*p_DMN_*_−*others*_ *<* 0.001, *p_V.Att_*_−*others*_ *<* 0.001; FDR-corrected). DMN and V.Att. did not differ in CNT or MCS; in UWS, DMN was less diverse than V.Att. Across groups, the limbic network displayed reduced diversity in MCS and UWS relative to CNT, and the DMN showed reduced diversity in UWS relative to both CNT and MCS.

### Local level of non-equilibrium

Node-wise analysis of local FDT deviation identified 59, 89 and 54/90 regions differing significantly in the CNT-MCS, CNT-UWS, and MCS-UWS comparisons, respectively (Figure 4.a; FDR-corrected). All significant t-values were positive, indicating greater downstream hierarchical influence (from the node to the whole-brain system) in the first group of each comparison. These local effects underlie the global and RSN-level differences. The extent to which local differences concentrate within each RSN was further examinated via the RSN’s affected volume fraction.

The largest CNT-MCS differences (Table 2) were found in bilateral supplementary motor area, median cingulate and paracingulate gyri, insula, inferior temporal gyrus, anterior cingulate and paracingulate gyri, temporal pole (superior temporal gyrus), paracentral lobule, and superior frontal gyrus (dorsolateral); and right thalamus and putamen (Figure 4.a, left). Overall, regions with significantly reduced FDT deviation in MCS relative to controls most strongly involved the somatomotor dorsal attention ventral attention default mode networks (Figure 4.a, left). Within the homogeneous flattening of local deviation observed in UWS relative to controls (Table 3), the largest differences were observed in bilateral superior frontal gyrus (dorsolateral and medial), supplementary motor area, (median and anterior) cingulate and paracingulate gyri, paracentral lobule, hippocampus, precentral gyrus, and middle frontal gyrus; left temporal pole (superior temporal gyrus), inferior temporal gyrus, middle occipital gyrus, and precuneus; and right putamen, thalamus, insula, and superior parietal gyrus. All networks were close to fully involved in these differences. For MCS-UWS (Table 4), the largest local differences were found in bilateral superior frontal gyrus (medial and dorsolateral), cuneus, lingual gyrus, calcarine fissure; left precentral gyrus, postcentral gyrus, olfactory cortex, and supramarginal gyrus; and right pallidum, superior occipital gyrus, and thalamus. The visual control dorsal attention and somatomotor networks showed the largest affected volume fractions.

**Table 2:** Regions with significant local FDT deviation differences: CNT vs MCS. Positive *t* indicates greater values in CNT.

| Label | Region | $t$-value |
| --- | --- | --- |
| 10 | L Supplementary Motor Area | 3.3471 |
| 17 | L Median Cingulate and Paracingulate Gyri | 3.2912 |
| 56 | R Paracentral Lobule | 3.2236 |
| 15 | L Insula | 3.1870 |
| 42 | L Temporal Pole: Superior Temporal Gyrus | 3.0399 |
| 76 | R Insula | 2.9876 |
| 45 | L Inferior Temporal Gyrus | 2.9099 |
| 52 | R Thalamus | 2.9067 |
| 75 | R Anterior Cingulate and Paracingulate Gyri | 2.8891 |
| 74 | R Median Cingulate and Paracingulate Gyri | 2.8838 |
| 89 | R Superior Frontal Gyrus, Dorsolateral | 2.8565 |
| 54 | R Lenticular Nucleus, Putamen | 2.8537 |
| 81 | R Supplementary Motor Area | 2.8484 |
| 16 | L Anterior Cingulate and Paracingulate Gyri | 2.7087 |
| 62 | R Postcentral Gyrus | 2.6799 |
| 61 | R Superior Parietal Gyrus | 2.6770 |
| 46 | R Inferior Temporal Gyrus | 2.6722 |
| 87 | R Middle Frontal Gyrus | 2.6643 |
| 34 | L Precuneus | 2.6273 |
| 35 | L Paracentral Lobule | 2.6188 |
| 90 | R Precentral Gyrus | 2.5680 |
| 12 | L Superior Frontal Gyrus, Medial | 2.5629 |
| 72 | R Hippocampus | 2.5544 |
| 63 | R Fusiform Gyrus | 2.5329 |
| 49 | R Temporal Pole: Superior Temporal Gyrus | 2.5310 |
| 64 | R Inferior Occipital Gyrus | 2.4613 |
| 19 | L Hippocampus | 2.4593 |
| 28 | L Fusiform Gyrus | 2.4464 |
| 2 | L Superior Frontal Gyrus, Dorsolateral | 2.4438 |
| 26 | L Middle Occipital Gyrus | 2.4416 |
| *Continues on next page* |  |  |

| Label | Region | *t*-value |
| --- | --- | --- |
| 65 | R Middle Occipital Gyrus | 2.4403 |
| 43 | L Middle Temporal Gyrus | 2.4238 |
| 57 | R Precuneus | 2.4201 |
| 53 | R Lenticular Nucleus, Pallidum | 2.3003 |
| 21 | L Amygdala | 2.2613 |
| 79 | R Superior Frontal Gyrus, Medial | 2.2196 |
| 55 | R Caudate Nucleus | 2.2133 |
| 30 | L Superior Parietal Gyrus | 2.1567 |
| 39 | L Thalamus | 2.1504 |
| 1 | L Precentral Gyrus | 2.1088 |
| 82 | R Rolandic Operculum | 2.0958 |
| 18 | L Posterior Cingulate Gyrus | 2.0818 |
| 4 | L Middle Frontal Gyrus | 2.0428 |
| 41 | L Superior Temporal Gyrus | 2.0332 |
| 27 | L Inferior Occipital Gyrus | 2.0147 |
| 20 | L Parahippocampal Gyrus | 2.0118 |
| 25 | L Superior Occipital Gyrus | 1.9715 |
| 71 | R Parahippocampal Gyrus | 1.9455 |
| 73 | R Posterior Cingulate Gyrus | 1.9369 |
| 40 | L Heschl Gyrus | 1.8853 |
| 37 | L Lenticular Nucleus, Putamen | 1.8469 |
| 9 | L Rolandic Operculum | 1.8365 |
| 36 | L Caudate Nucleus | 1.8180 |
| 44 | L Temporal Pole: Middle Temporal Gyrus | 1.8078 |
| 31 | L Inferior Parietal Gyri | 1.7813 |
| 70 | R Amygdala | 1.7659 |
| 13 | L Superior Frontal Gyrus, Medial Orbital | 1.7522 |
| 78 | R Superior Frontal Gyrus, Medial Orbital | 1.7364 |
| 47 | R Temporal Pole: Middle Temporal Gyrus | 1.6887 |

**Table 3:**
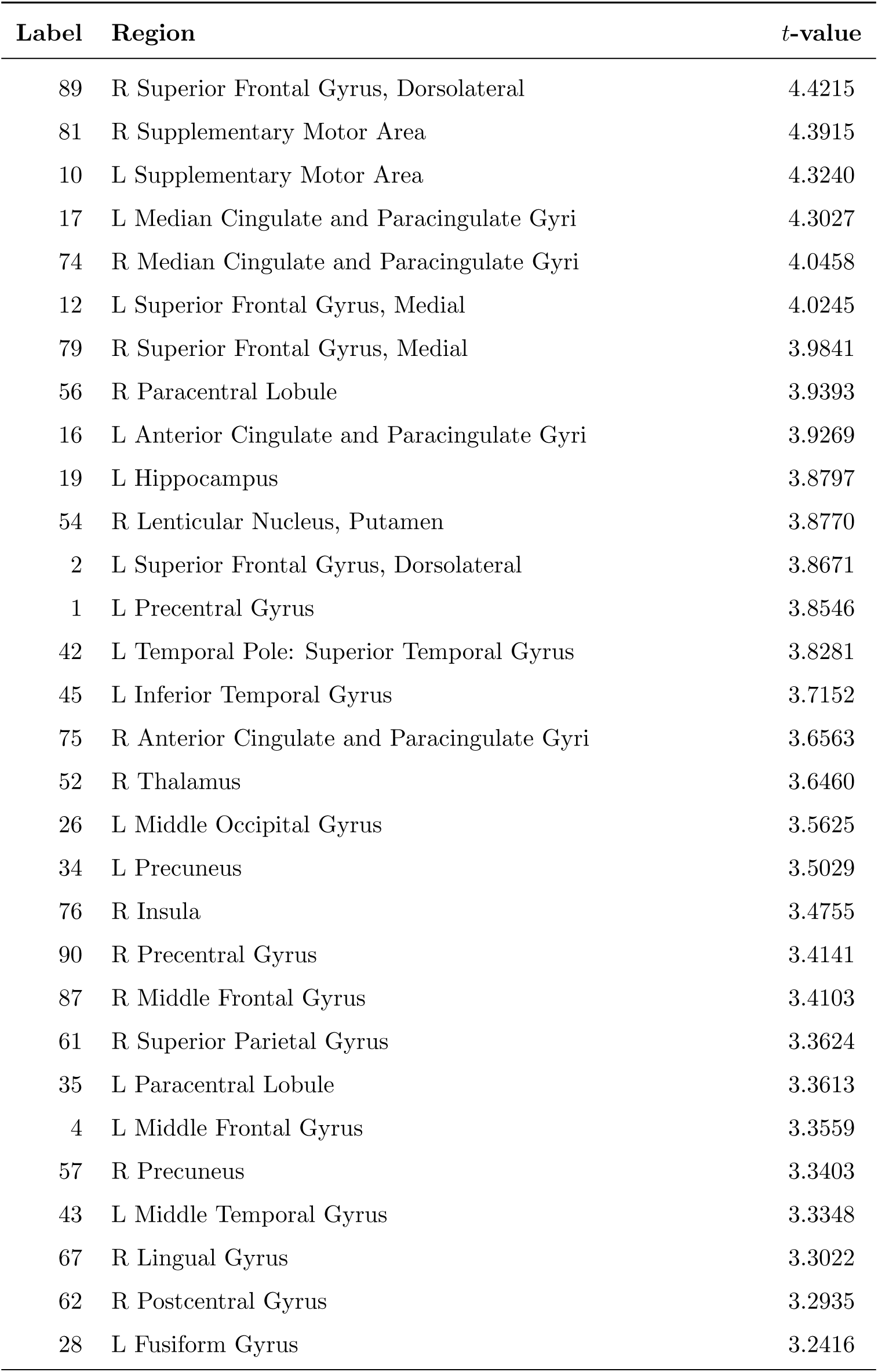
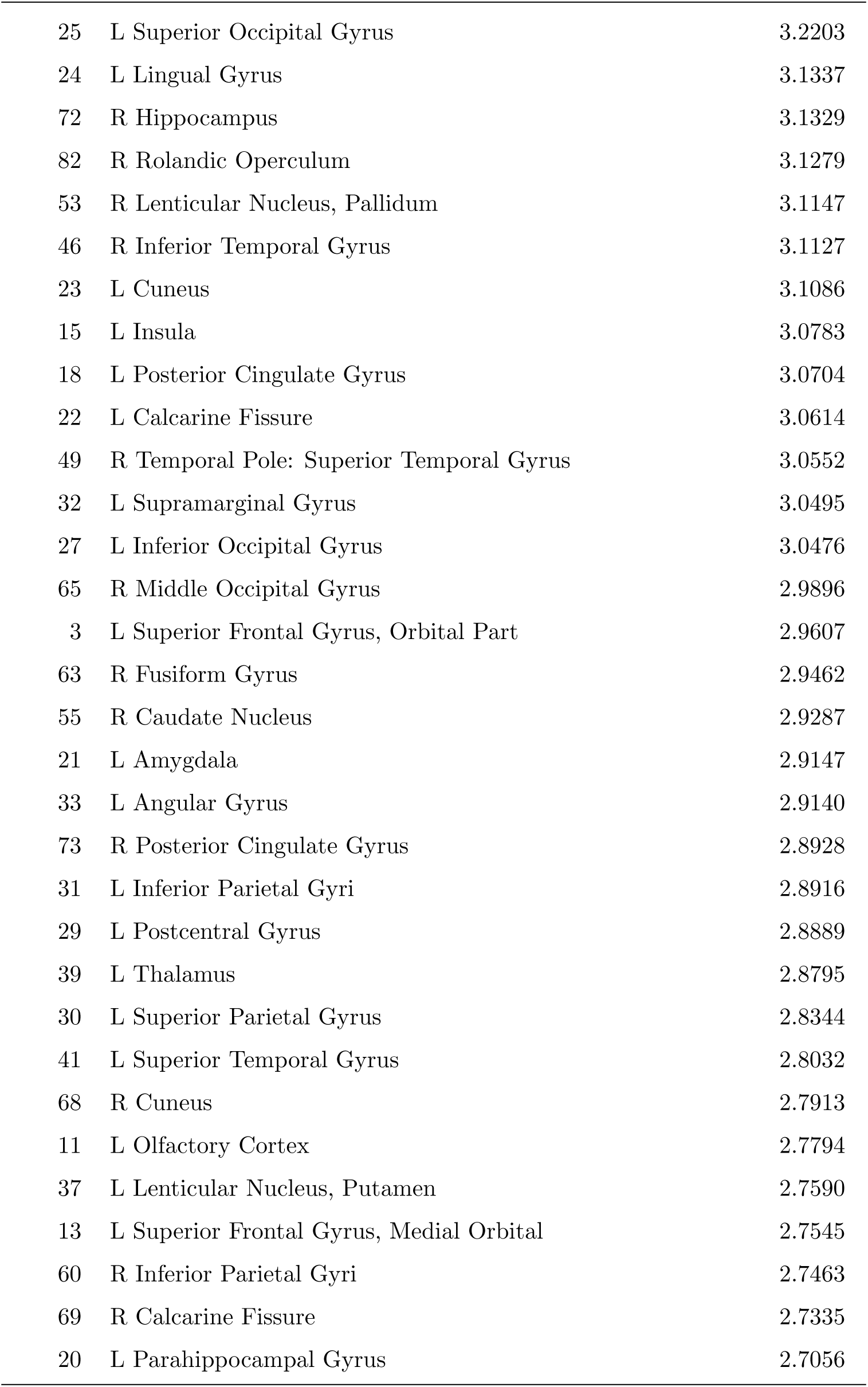
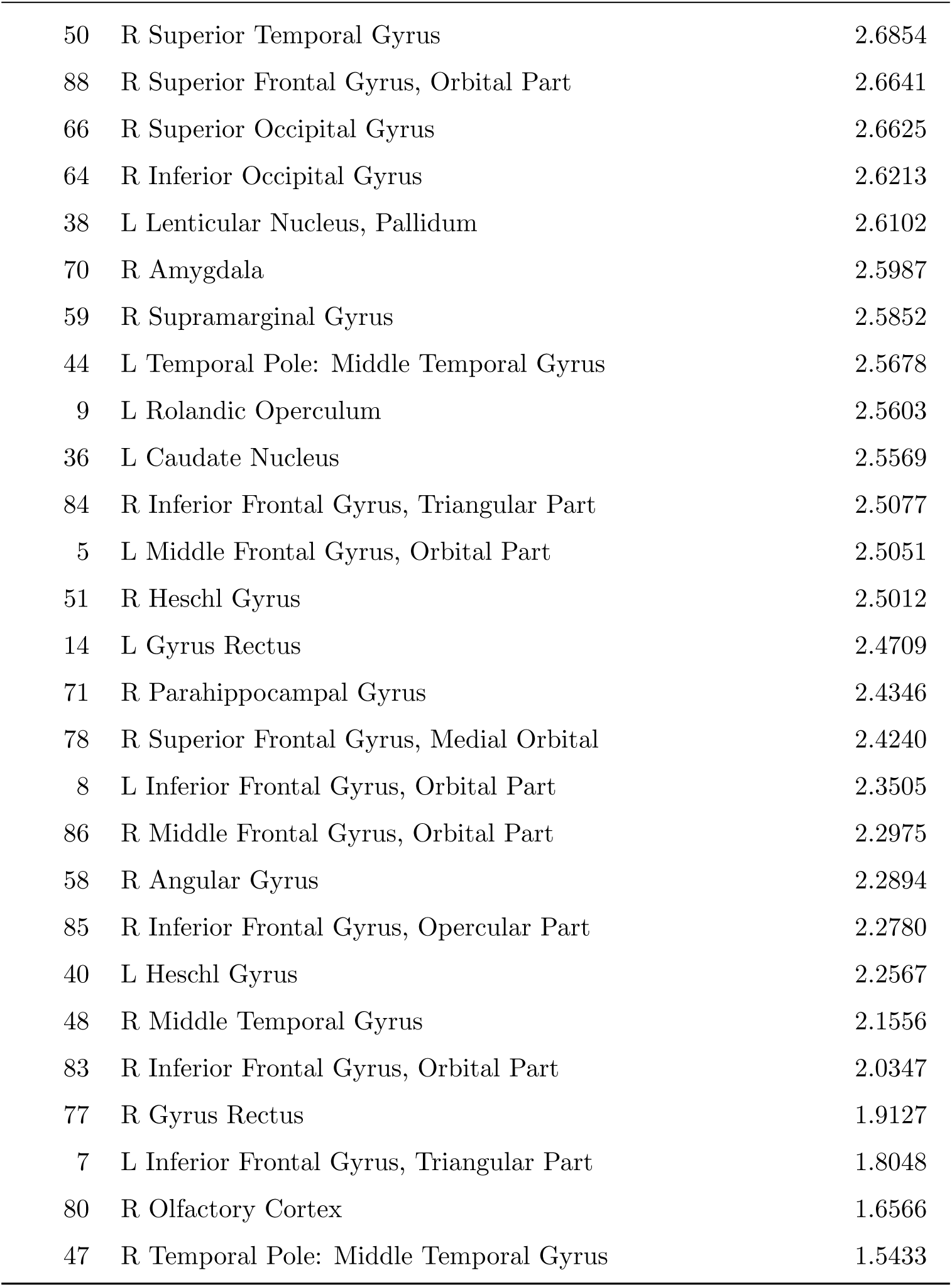
Regions with significant local FDT deviation differences: CNT vs UWS. Positive *t* indicates greater values in CNT.

| Label | Region | $t$-value |
| --- | --- | --- |
| 89 | R Superior Frontal Gyrus, Dorsolateral | 4.4215 |
| 81 | R Supplementary Motor Area | 4.3915 |
| 10 | L Supplementary Motor Area | 4.3240 |
| 17 | L Median Cingulate and Paracingulate Gyri | 4.3027 |
| 74 | R Median Cingulate and Paracingulate Gyri | 4.0458 |
| 12 | L Superior Frontal Gyrus, Medial | 4.0245 |
| 79 | R Superior Frontal Gyrus, Medial | 3.9841 |
| 56 | R Paracentral Lobule | 3.9393 |
| 16 | L Anterior Cingulate and Paracingulate Gyri | 3.9269 |
| 19 | L Hippocampus | 3.8797 |
| 54 | R Lenticular Nucleus, Putamen | 3.8770 |
| 2 | L Superior Frontal Gyrus, Dorsolateral | 3.8671 |
| 1 | L Precentral Gyrus | 3.8546 |
| 42 | L Temporal Pole: Superior Temporal Gyrus | 3.8281 |
| 45 | L Inferior Temporal Gyrus | 3.7152 |
| 75 | R Anterior Cingulate and Paracingulate Gyri | 3.6563 |
| 52 | R Thalamus | 3.6460 |
| 26 | L Middle Occipital Gyrus | 3.5625 |
| 34 | L Precuneus | 3.5029 |
| 76 | R Insula | 3.4755 |
| 90 | R Precentral Gyrus | 3.4141 |
| 87 | R Middle Frontal Gyrus | 3.4103 |
| 61 | R Superior Parietal Gyrus | 3.3624 |
| 35 | L Paracentral Lobule | 3.3613 |
| 4 | L Middle Frontal Gyrus | 3.3559 |
| 57 | R Precuneus | 3.3403 |
| 43 | L Middle Temporal Gyrus | 3.3348 |
| 67 | R Lingual Gyrus | 3.3022 |
| 62 | R Postcentral Gyrus | 3.2935 |
| 28 | L Fusiform Gyrus | 3.2416 |
| *Continues on next page* |  |  |

| Label | Region | *t*-value |
| --- | --- | --- |
| 25 | L Superior Occipital Gyrus | 3.2203 |
| 24 | L Lingual Gyrus | 3.1337 |
| 72 | R Hippocampus | 3.1329 |
| 82 | R Rolandic Operculum | 3.1279 |
| 53 | R Lenticular Nucleus, Pallidum | 3.1147 |
| 46 | R Inferior Temporal Gyrus | 3.1127 |
| 23 | L Cuneus | 3.1086 |
| 15 | L Insula | 3.0783 |
| 18 | L Posterior Cingulate Gyrus | 3.0704 |
| 22 | L Calcarine Fissure | 3.0614 |
| 49 | R Temporal Pole: Superior Temporal Gyrus | 3.0552 |
| 32 | L Supramarginal Gyrus | 3.0495 |
| 27 | L Inferior Occipital Gyrus | 3.0476 |
| 65 | R Middle Occipital Gyrus | 2.9896 |
| 3 | L Superior Frontal Gyrus, Orbital Part | 2.9607 |
| 63 | R Fusiform Gyrus | 2.9462 |
| 55 | R Caudate Nucleus | 2.9287 |
| 21 | L Amygdala | 2.9147 |
| 33 | L Angular Gyrus | 2.9140 |
| 73 | R Posterior Cingulate Gyrus | 2.8928 |
| 31 | L Inferior Parietal Gyri | 2.8916 |
| 29 | L Postcentral Gyrus | 2.8889 |
| 39 | L Thalamus | 2.8795 |
| 30 | L Superior Parietal Gyrus | 2.8344 |
| 41 | L Superior Temporal Gyrus | 2.8032 |
| 68 | R Cuneus | 2.7913 |
| 11 | L Olfactory Cortex | 2.7794 |
| 37 | L Lenticular Nucleus, Putamen | 2.7590 |
| 13 | L Superior Frontal Gyrus, Medial Orbital | 2.7545 |
| 60 | R Inferior Parietal Gyri | 2.7463 |
| 69 | R Calcarine Fissure | 2.7335 |
| 20 | L Parahippocampal Gyrus | 2.7056 |

| Label | Region | *t*-value |
| --- | --- | --- |
| 50 | R Superior Temporal Gyrus | 2.6854 |
| 88 | R Superior Frontal Gyrus, Orbital Part | 2.6641 |
| 66 | R Superior Occipital Gyrus | 2.6625 |
| 64 | R Inferior Occipital Gyrus | 2.6213 |
| 38 | L Lenticular Nucleus, Pallidum | 2.6102 |
| 70 | R Amygdala | 2.5987 |
| 59 | R Supramarginal Gyrus | 2.5852 |
| 44 | L Temporal Pole: Middle Temporal Gyrus | 2.5678 |
| 9 | L Rolandic Operculum | 2.5603 |
| 36 | L Caudate Nucleus | 2.5569 |
| 84 | R Inferior Frontal Gyrus, Triangular Part | 2.5077 |
| 5 | L Middle Frontal Gyrus, Orbital Part | 2.5051 |
| 51 | R Heschl Gyrus | 2.5012 |
| 14 | L Gyrus Rectus | 2.4709 |
| 71 | R Parahippocampal Gyrus | 2.4346 |
| 78 | R Superior Frontal Gyrus, Medial Orbital | 2.4240 |
| 8 | L Inferior Frontal Gyrus, Orbital Part | 2.3505 |
| 86 | R Middle Frontal Gyrus, Orbital Part | 2.2975 |
| 58 | R Angular Gyrus | 2.2894 |
| 85 | R Inferior Frontal Gyrus, Opercular Part | 2.2780 |
| 40 | L Heschl Gyrus | 2.2567 |
| 48 | R Middle Temporal Gyrus | 2.1556 |
| 83 | R Inferior Frontal Gyrus, Orbital Part | 2.0347 |
| 77 | R Gyrus Rectus | 1.9127 |
| 7 | L Inferior Frontal Gyrus, Triangular Part | 1.8048 |
| 80 | R Olfactory Cortex | 1.6566 |
| 47 | R Temporal Pole: Middle Temporal Gyrus | 1.5433 |

**Table 4:** Regions with significant local FDT deviation differences: MCS vs UWS. Positive *t* indicates greater values in MCS.

| Label | Region | $t$-value |
| --- | --- | --- |
| 1 | L Precentral Gyrus | 3.5982 |
| 29 | L Postcentral Gyrus | 3.5839 |
| 79 | R Superior Frontal Gyrus, Medial | 3.5530 |
| 12 | L Superior Frontal Gyrus, Medial | 3.5286 |
| 89 | R Superior Frontal Gyrus, Dorsolateral | 3.4468 |
| 68 | R Cuneus | 3.1279 |
| 23 | L Cuneus | 3.0961 |
| 53 | R Lenticular Nucleus, Pallidum | 3.0337 |
| 24 | L Lingual Gyrus | 2.9429 |
| 11 | L Olfactory Cortex | 2.9399 |
| 22 | L Calcarine Fissure | 2.9359 |
| 66 | R Superior Occipital Gyrus | 2.9117 |
| 2 | L Superior Frontal Gyrus, Dorsolateral | 2.8798 |
| 32 | L Supramarginal Gyrus | 2.8709 |
| 52 | R Thalamus | 2.8599 |
| 4 | L Middle Frontal Gyrus | 2.7887 |
| 16 | L Anterior Cingulate and Paracingulate Gyri | 2.7366 |
| 81 | R Supplementary Motor Area | 2.7079 |
| 43 | L Middle Temporal Gyrus | 2.6245 |
| 67 | R Lingual Gyrus | 2.5411 |
| 19 | L Hippocampus | 2.5191 |
| 7 | L Inferior Frontal Gyrus, Triangular Part | 2.5094 |
| 69 | R Calcarine Fissure | 2.5059 |
| 6 | L Inferior Frontal Gyrus, Opercular Part | 2.4993 |
| 28 | L Fusiform Gyrus | 2.4826 |
| 18 | L Posterior Cingulate Gyrus | 2.4712 |
| 14 | L Gyrus Rectus | 2.4417 |
| 26 | L Middle Occipital Gyrus | 2.4202 |
| 38 | L Lenticular Nucleus, Pallidum | 2.3900 |
| 84 | R Inferior Frontal Gyrus, Triangular Part | 2.3737 |
| *Continues on next page* |  |  |

| Label | Region | *t*-value |
| --- | --- | --- |
| 41 | L Superior Temporal Gyrus | 2.3684 |
| 39 | L Thalamus | 2.3564 |
| 10 | L Supplementary Motor Area | 2.3216 |
| 50 | R Superior Temporal Gyrus | 2.2825 |
| 73 | R Posterior Cingulate Gyrus | 2.2420 |
| 25 | L Superior Occipital Gyrus | 2.2400 |
| 55 | R Caudate Nucleus | 2.2081 |
| 82 | R Rolandic Operculum | 2.1206 |
| 51 | R Heschl Gyrus | 2.0915 |
| 77 | R Gyrus Rectus | 2.0796 |
| 56 | R Paracentral Lobule | 2.0570 |
| 70 | R Amygdala | 2.0478 |
| 17 | L Median Cingulate and Paracingulate Gyri | 2.0328 |
| 60 | R Inferior Parietal Gyri | 2.0259 |
| 33 | L Angular Gyrus | 2.0223 |
| 34 | L Precuneus | 1.9987 |
| 61 | R Superior Parietal Gyrus | 1.9837 |
| 54 | R Lenticular Nucleus, Putamen | 1.9694 |
| 42 | L Temporal Pole: Superior Temporal Gyrus | 1.9669 |
| 57 | R Precuneus | 1.9587 |
| 75 | R Anterior Cingulate and Paracingulate Gyri | 1.9031 |
| 3 | L Superior Frontal Gyrus, Orbital Part | 1.8575 |
| 86 | R Middle Frontal Gyrus, Orbital Part | 1.8328 |
| 30 | L Superior Parietal Gyrus | 1.7584 |

Three subsets of brain regions –defined by significance patterns across pairwise comparisons– were of particular interest (Figure 5.a). Case 1 comprised areas with a graded decrease in deviation across CNT, MCS and UWS (32/90). Case 2 included regions with reduced deviation in UWS relative to both CNT and MCS, with no difference between the latter (21/90). These two subsets primarily account for the global and RSN differences between the DoC groups. Case 3 comprised regions with similarly reduced deviation in both DoC relative to CNT (27/90), potentially reflecting trains shared across DoC that cannot be inferred from global or RSN averages alone. The RSN volume fraction covered by each subset was computed as for the pairwise analyses (Figure 5.b). For completeness, that 10 regions did not fall into any of these subsets. Accordingly, network fractions across cases do not sum to one.

**Figure 5.**
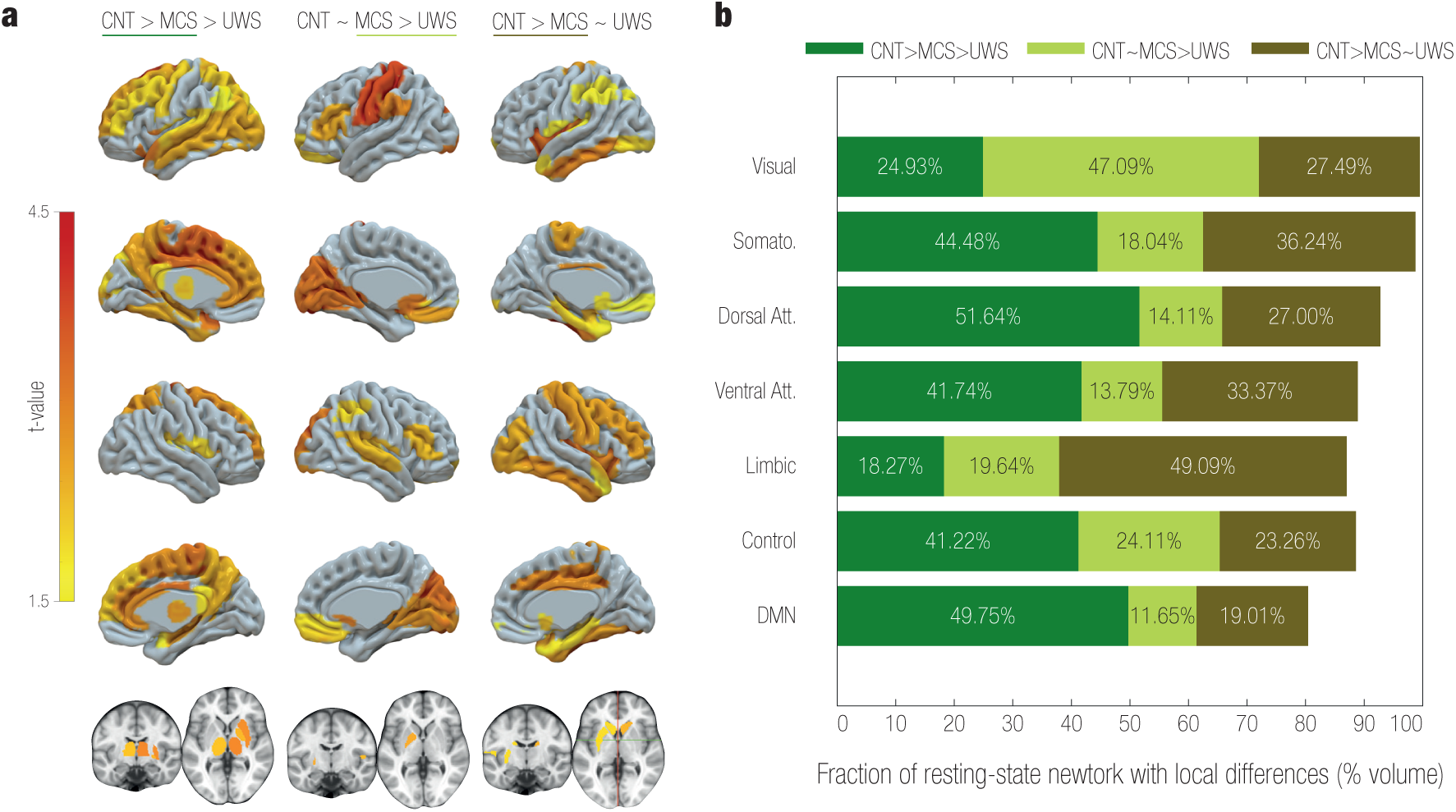
Local FDT deviation across comparisons. a) Cases of interest: regions with a gradual decrease in local deviation across groups, local differences and local similarities between MCS and UWS regarding controls. Among local differences observed in pairwise group comparisons (Figure 4.a), 80/90 regions exhibited three distinct patterns of significance across the comparisons, corresponding to three cases of interest. Case 1 (left, CNT *>* MCS *>* UWS): 32 regions showed a gradual decrease of local non-equilibrium across the three groups. Case 2 (middle, CNT = MCS *>* UWS): 21 regions differed in control-UWS and MCS-UWS comparisons, but not between controls and MCS. Case 3 (right, CNT *>* MCS = UWS): 27 regions differed in both control-DoC comparisons, but not between MCS and UWS, indicating similarly reduced FDT deviation for MCS and UWS regarding controls. For each case, the subsets of regions are coloured according to the t-values of the underscored pair of groups. Gray areas did not belong to the case’s subset. **b) Fraction of resting-state networks involved in the cases.** For the cases in a), summarized in the legend, the volume fraction of network occupied by each subset of regions was calculated. The default mode network was particularly and largely involved in gradual closer-to-equilibrium dynamics with decreasing levels of consciousness (case 1). Similarly, the dorsal attention network mostly decreased gradually, but was also prominently engaged in local similarities between DoC (case 3). Conversely, the limbic network was mostly related to similarities between MCS and UWS regarding controls (case 3). Last, the visual network showed the greatest fraction of differences between MCS and UWS related to control-like local deviations in MCS (case 2).

Case 1 regions –progressively closer to equilibrium with decreasing levels of consciousness– occupied 51.64% of the dorsal attention, 49.75% of the default mode, 44.48% of the somatomotor, 41.74% of the ventral attention, 41.22% of the control, 24.93% of the visual, and 18.27% of the limbic networks. Among these, the largest MCS-UWS difference were found in bilateral superior frontal gyrus (medial and dorsolateral); left precentral gyrus, middle frontal gyrus, and anterior cingulate and paracingulate gyri; and right pallidum and thalamus. The largest CNT-MCS within Case 1 also included the bilateral supplementary motor area; left median cingulate and paracingulate gyri, temporal pole (superior temporal gyrus), and precuneus; and right paracentral lobule, and putamen.

Case 2 regions –reduced in UWS relative to both CNT and MCS, but not differing between the latter– covered 47.09% of the visual, 29.64% of the limbic, 24.11% of the control, 18.04% of the somatomotor, 14.11% of the dorsal attention, 13.79% of the ventral attention, and 11.65% of the default mode networks. The largest t-values in the MCS-UWS comparison within this subset were observed in bilateral cuneus and lingual gyrus; left postcentral gyrus, olfactory cortex, and supramarginal gyrus; and right superior occipital gyrus.

Case 3 –differing only in control-DoC comparisons– comprised 49.09% of the limbic, 36.24% of the somatomotor, 33.37% of the ventral attention, 27.49% of the visual, 27.00% of the dorsal attention, 23.26% of the control, and 19.01% of the default mode networks. The largest CNT-MCS differences within this subset were found in bilateral insula, and inferior temporal gyrus; left paracentral lobule; and right median cingulate and paracingulate gyri, postcentral gyrus, middle frontal gyrus, precentral gyrus, hippocampus, fusiform gyrus, temporal pole (superior temporal gyrus), inferior occipital gyrus, and middle occipital gyrus .

Among all networks, the DMN showed a distinctive pattern, being predominantly and extensively involved in Case 1 (i.e., exhibiting a gradual shift toward equilibrium with decreasing consciousness). The dorsal attention network also aligned with Case 1 but with less specificity (substantial fractions were involved in Cases 2 and 3). By contrast, the visual and limbic networks were more strongly associated with potential differences and similarities between the two DoC groups (Cases 2 and 3, respectively).

## Discussion

The functional hierarchical organization was assessed for controls (CNT), minimally conscious state (MCS), and unresponsive wakefulness state (UWS) using the model-based fluctuation-dissipation framework [41, 54]. FDT deviation is a perturbative measure of non-equilibrium interactions (asymmetric information flow) between brain regions. Global and resting-state network FDT deviation decreased stepwise with reduced levels of consciousness. Together with prior evidence for closer-to-equilibrium dynamics [44, 54] and reduced irreversibility [49, 52, 74] in states of reduced consciousness, our results position non-equilibrium dynamics as a signature of conscious capacity.

In non-equilibrium network dynamics, time-asymmetry in information flows relates to asymmetric generative effective coupling [41, 51, 53, 54]. In line with previous evidence, individualized whole-brain Hopf models shows less asymmetric generative effective coupling in DoC (Supplementary, Figure 7). However, by computing closed-form linear responses, our approach estimates how each region causally drives the rest of the system. The observed reduction in FDT deviation in DoC therefore indicates weakened asymmetric interactions even between nodes lacking direct anatomical connection, providing a principled form of functional hierarchy beyond asymmetries in generative effective coupling. Whereas temporal irreversibility has been used to quantify two-way-averaged asymmetry of ongoing activity [50, 52], here we report site-specific hierarchical drive over the whole system. The parallel drop in global hierarchical diversity indicates a collapse toward more uniform working points, aligning with prior reports of homogenized dynamics in DoC [24]. Furthermore, hierarchical diversity within RSNs was an order of magnitude lower than diversity across the whole brain, suggesting that network components, beyond being functionally coupled [67], behave more uniformly in their hierarchical interactions with the system. Across all groups, the default mode and ventral attention networks exhibited strikingly uniform hierarchy levels among their components. Overall, the results are consistent with theories proposing that consciousness requires directed integration across a functional hierarchy [30, 35].

**Figure 6.**
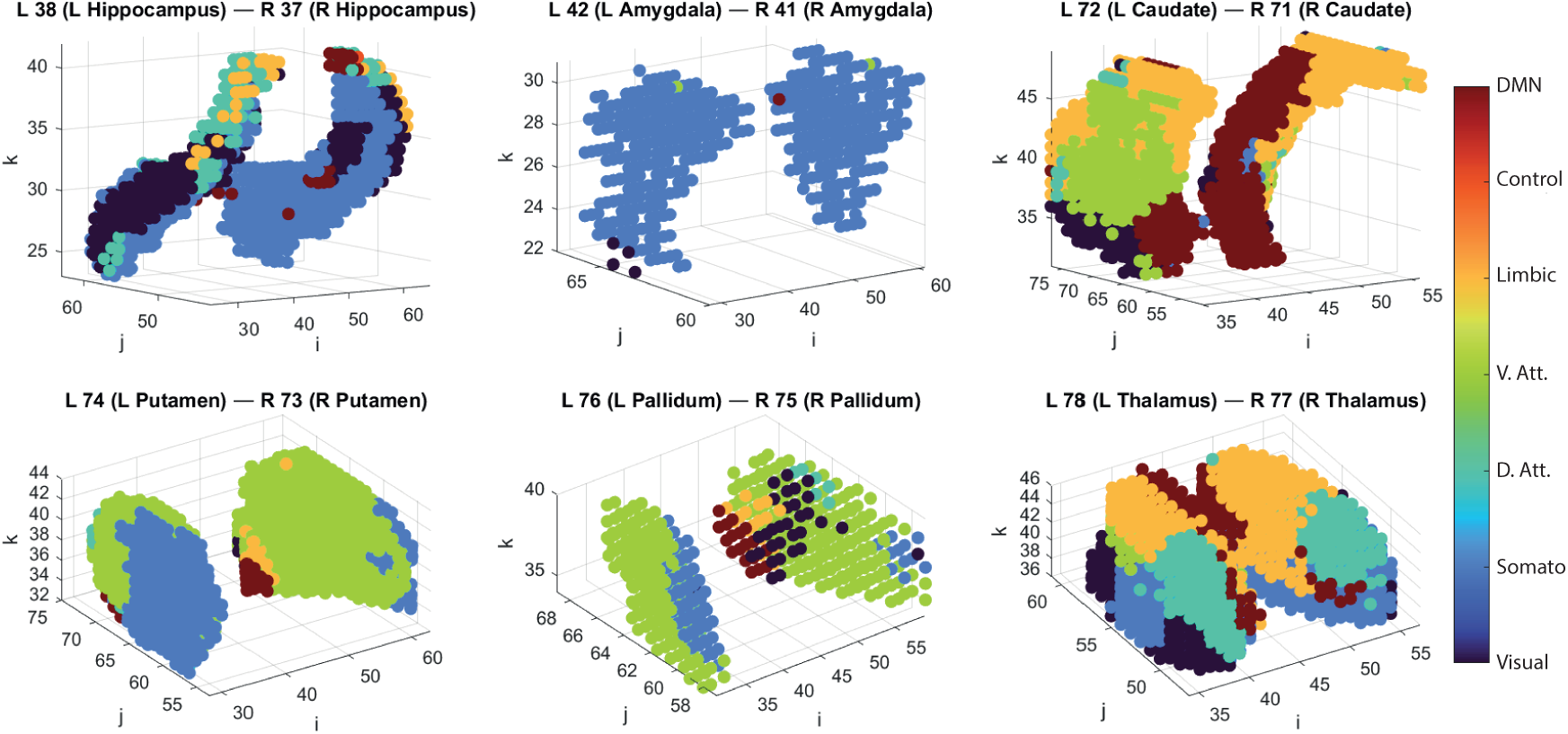
Subcortical-to-RSN assignments (Yeo 7). Each panel shows a homotopic AAL subcortical pair (L/R; 2mm voxels) in voxel coordinates (*i, j, k*), coloured by the assigned resting-state network. Assignments were obtained by computing subcortical–cortical functional connectivity in 100 unrelated HCP subjects, averaging correlations within each RSN, and labelling each subcortical voxel by the RSN with the maximal mean correlation. See Methods §.

**Figure 7.**
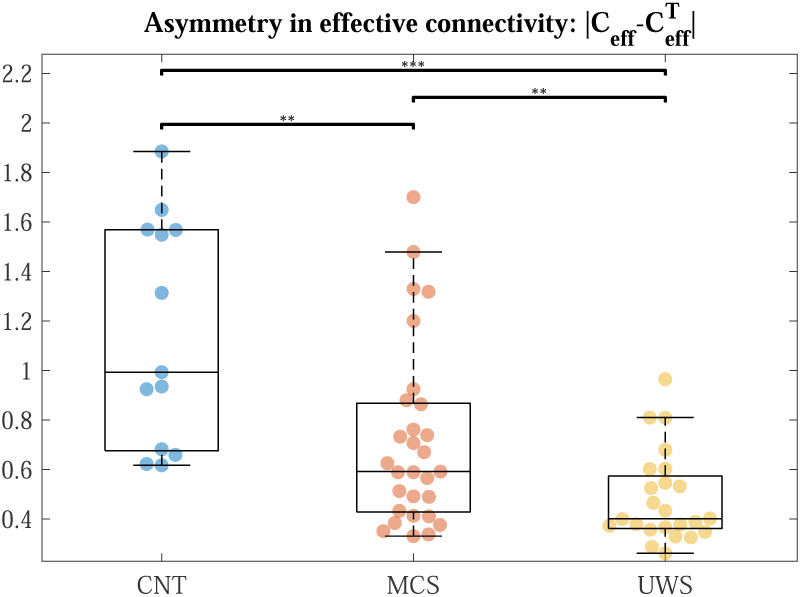
Asymmetry in generative effective connectivity. By fitting individualized whole-brain Hopf models to empirical data, we obtain the generative effective connectivities, *C_eff_* necessary for computing closed-form linear response to weak perturbations. We compute asymmetry of these matrices as |*C_eff_* − *C^T^* |.

Our findings complement perturbation-based indices such as PCI, which captures the spatiotemporal spread and algorithmic complexity of evoked responses [4, 37, 39]. Whereas PCI often yields similar values across a tested subset of targets, exhaustive exploration of perturbation sites reveals varying local FDT deviation values. Recent perturbation studies likewise report target-dependent responses [75, 76], suggesting that FDT deviation may be target-sensitive even for linear responses after infinitesimal drive, in contrast to over-threshold evoked responses typically used for PCI. In addition, whole-brain computational models enable exhaustive *in silico* exploration of perturbation sites and parameters, naturally integrating frameworks for causal probing and therapy design [26, 77].

Altered hierarchical level in regions within the default mode network was particularly associated with gradual closer-to-equilibrium interactions as consciousness decreased (Case 1, CNT *>* MCS *>* UWS). Prior work links DMN activity with conscious awareness [78–80], with DMN connectivity distinguishing between levels of consciousness [8, 52, 81–83], and correlating with the CRS-R score [11, 84, 85]. Our FDT-based hierarchy readout extends this literature by showing that, gradually across MCS and UWS, DMN component’s loose their ability to causally influence the system. Regions in the dorsal attention network (DAT) were also conspicuously related to this outcome. Consistent with this graded DMN–DAT hierarchy loss, frontal/posterior HD-tDCS increases long-range anterior–posterior transfers and DMN activation [75], and TMS–EEG shows markedly lower fronto–parietal bidirectional flow in UWS than MCS, with clinical correlations [76]. Stepwise reductions in hierarchical level were also prominent in the thalamus and basal ganglia (pallidum and putamen). Disrupted thalamo-cortical interplay is a hallmark of DoC [24, 86–88], and thalamic activity is known to diminish with anaesthesia [89], with connectivity varying across sedation levels [90]. Thalamic stimulation during anesthesia elicits arousal in non-human primates [27, 28], and has motivated the development of computational models that provide mechanistic explanations while enabling low-risk exploration [91].

The visual network was the most engaged by regions with recovered hierarchical level in MCS but reduced in UWS (Case 2, CNT ≈ MCS *>* UWS). Visual system connectivity has shown strong MCS-UWS discriminative capacity in prior work [85, 92]. Recovery of hierarchical levels in the visual network in MCS suggests that exogenous sensory drive can still propagate causally to the broader system, in line with activation of higher-order visual areas in MCS but not UWS [93]. Because activity in visual cortex does not guarantee conscious perception [94, 95], these findings reinforce the need to probe neural correlates of conscious visual perception alongside behavioural signs such as pursuit/fixation [96, 97]. In addition, the largest MCS-UWS difference was found in the left postcentral gyrus, which is known to preserve functional coupling in MCS but not UWS [98].

Regions with similar hierarchy loss across DoC (Case 3, CNT *>* MCS ≈ UWS) highly impacted the limbic network, with a clear paralimbic and anterior-temporal emphasis. This finding dovetails with reports that limbic circuitry is the only network exhibiting similar non-equilibrium loss in MCS and UWS under bidirectional averaging [52]. Connectivity studies have further identified gradual hyperconnectivity between limbic regions and the DMN with decreasing consciousness in DoC [99]. Notably, bilateral insula showed the largest differences relative to controls, aligning with its role in DMN–DAT switching and perceptual awareness [100].

The FDT framework offers a computationally efficient, model-based non-equlibrium hierarchy readout that separates DoC groups across scales and can be operationalised for clinical stratification, intervention planning and monitoring. Because it is tied to directed interactions, it also provides falsifiable predictions that align with empirical perturbation studies (e.g., larger gains in MCS than UWS at Case 1 and 2 loci [75, 76, 101]) and suggests putative targets for neuromodulation . Our findings were observed irrespective of disorder aetiology, providing insights into common traits of each condition. However, these inferences are cross-sectional and model-dependent. Future work should replicate across parcellations and adjust for aetiology, lesion burden, and medication. The use of individual anatomical connectivities along with node-wise optimization of local dynamics should yield more accurate predictions. The RSN analysis would benefit from one-to-one node–network mappings and from testing sensitivity to the subcortical-to-cortical assignment derived from HCP data. Longitudinal sampling will be essential to establish prognostic value and to determine recovery-related shifts in non-equilibrium hierarchy toward control-like levels. Finally, prospective studies should screen stimulation sites and test whether targeted stimulation increases global/RSN averages for advancing patient-specific stimulation strategies [4, 26, 77].

Overall, the FDT framework captures differential reductions of non-equilibrium across DoC, associated with a breakdown of hierarchical organization. By uniting generative modelling with a principled non-equilibrium measure, we isolate directional deficits in large-scale hierarchical interactions, clarifying how hierarchical drive from DMN/DAT and thalamo–basal-ganglia hubs degrades with decreasing consciousness, distinguishing potential recovery-linked differences between MCS and UWS among visual territories, and revealing limbic alterations shared across DoC. These findings advance a model-based account of how conscious capacity diminishes as brain dynamics approach equilibrium and motivate testable neuromodulation targets for future studies.

## Acknowledgments

The authors express gratitude to all the individuals who participated in the studies. We are as well grateful to the patient’s families, whose consent and understanding make advancements in the field possible. We extend our gratitude to the clinicians and personnel at Ĥopital de la Pitié Salp^etrìere (Paris, France) for their dedication and support. M.M. is a PRE fellow (PRE2022-101417) supported by the Grant CEX2021-001195-M-20-5 co-funded by EU and the Spanish ”Ministerio de Ciencia, Innovación y Universidades”. A.E. was supported by the project eBRAIN-Health—Actionable Multilevel Health Data (id 101058516), funded by EU Horizon Europe and by the Grant PID2022-136216NBI00, funded by MICIU/AEI/10.13039/501100011033, and “ERDF A way of making Europe”, ERDF, EU. D.M. and J.D.S. were supported by funding from the EU ERAPerMed Joint Translational Call for Proposals for “Personalised Medicine: Multidisciplinary research towards implementation” (ERA PerMed JTC2019). J.D.S. was also supported by the FLAG-ERA JTC2021 project ModelDXConsciousness (Human Brain Project Partnering Project). Y.S.P.was supported by European Union’s Horizon 2020 research and innovation programme under the Marie Sklodowska-Curie grant 896354, and ‘ERDF A way of making Europe’, ERDF, EU, Project NEurological MEchanismS of Injury, and Sleep-like cellular dynamics (NEMESIS; ref. 101071900) funded by the EU ERC Synergy Horizon Europe. M.L.K. was supported by the Center for Music in the Brain, funded by the Danish National Research Foundation (DNRF117), and Centre for Eudaimonia and Human Flourishing at Linacre College funded by the Pettit and Carlsberg Foundations. G.D. was supported by grant no. PID2022-136216NB-I00 funded by MI-CIU/AEI/10.13039/501100011033 and by ‘ERDF A way of making Europe’, ERDF, EU, Project NEurological MEchanismS of Injury, and Sleep-like cellular dynamics (NEMESIS; ref. 101071900) funded by the EU ERC Synergy Horizon Europe, and AGAUR research support grant (ref. 2021 SGR 00917) funded by the Department of Research and Universities of the Generalitat of Catalunya. Data were provided [in part] by the Human Connectome Project, WU-Minn Consortium (Principal Investigators: David Van Essen and Kamil Ugurbil; 1U54MH091657) funded by the 16 NIH Institutes and Centers that support the NIH Blueprint for Neuroscience Research; and by the McDonnell Center for Systems Neuroscience at Washington University. Specifically, for associating subcortical voxels to cortical resting-state networks.

## Author contributions

Martínez-Marín M: Conceptualisation, Methodology, Software, Formal analysis, Visualization, Writing–Original Draft, Writing–Review & Editing. Vohryzek J: Conceptualisation, Methodology, Resources, Writing–Review & Editing. Escrichs A: Methodology, Resources, Writing–Review & Editing. Manasova D: Resources, Writing–Review. Sitt J.D.: Resources, Writing–Review. Kringelbach L.M.: Conceptualisation, Writing–Review & Editing. Sanz Perl Y: Conceptualisation, Methodology, Software, Supervision, Writing–Review & Editing Deco G: Conceptualisation, Funding acquisition, Methodology, Software, Supervision, Writing–Review & Editing

## Data availability

Due to the sensitive nature of clinical data, the Disorders of Consciousness dataset is available upon consultation with Société de Réanimation de Langue Française (https://www.srlf.org/, reference: M-neuro-Doc). The 100 unrelated subjects dataset used for subcortical-RSN mapping is freely available from HCP.

## Code availability

Code underlying the results in this paper is available at https://github.com/marian-martmar.

## Supplementary Material

